# The identification of temporal communities through trajectory clustering correlates with single-trial behavioural fluctuations in neuroimaging data

**DOI:** 10.1101/617027

**Authors:** William Hedley Thompson, Jessey Wright, James M. Shine, Russell A. Poldrack

## Abstract

Interacting sets of nodes and fluctuations in their interaction are important properties of a dynamic network system. In some cases the edges reflecting these interactions are directly quantifiable from the data collected. However, in many cases (such as functional magnetic resonance imaging (fMRI) data), the edges must be inferred from statistical relations between the nodes. Here we present a new method, Temporal Communities through Trajectory Clustering (TCTC), that derives time-varying communities directly from time-series data collected from the nodes in a network. First, we verify TCTC on resting and task fMRI data by showing that time-averaged results correspond with expected static connectivity results. We then show that the time-varying communities correlate and predict single-trial behaviour. This new perspective on temporal community detection of node-collected data identifies robust communities revealing ongoing spatiotemporal community configurations during task performance.

## Introduction

Many empirical phenomena can be mathematically described as networks, and an important property of networks is the presence of community structure. Communities are sets of nodes that are more strongly interconnected with one another compared to the rest of the network (Fortunato and Hric, 2016; Newman, 2010). When collecting network data, information is sampled from different nodes or edges. Edge-collected data, such as the number of emails sent between people, can seamlessly translate into a network representation. In contrast, node-collected data requires that edges be inferred based on a statistical relationship between nodes. This procedure is typical of most non-invasive neuroimaging techniques, where recorded brain regions have their connectivity inferred from statistical relationships between the nodes’ time-series (e.g. using Pearson’s correlation). Only after this inference step can different network properties be calculated.

Whereas early work focused on static network structures, there is increasing interest in identifying how networks change over time (Holme and Saramäki, 2012). In node-collected cases, edges must also be inferred over time (e.g. using sliding-window techniques), which involves a trade-off between temporal resolution and estimate edge precision. Using more time-points to assist the edge inference will decrease the temporal resolution of the network, whereas using fewer time-points will entail a less precise estimate of the edge due to the instability of statistical estimates based on small numbers of samples. One must thus choose between increasing uncertainty or losing temporal resolution, both of which amplify uncertainty in the inferred edges within the temporal network. This trade-off will distort and blur properties derived from the representation, such as community detection.

Temporal community detection identifies fluctuating communities over time and can quantify changes in the interaction or groupings of nodes (Bazzi et al., 2016; Mucha, Richardson, Macon, Porter, and Onnela, 2010; Peixoto and Rosvall, 2017; Rosvall and Bergstrom, 2010). Community detection algorithms also contain uncertainty, as alternative methods will often produce slightly different results. Given that well-established static community detection algorithms can give vastly different communities when applied to complex real-world networks with noise (Hric, Darst, and Fortunato, 2014), temporal extensions of these algorithms offer no inherent solution to uncertainty in the community detection step. Thus, two-step solutions to estimate communities from node-collected data (i.e. edge inference and then community detection) will propagate and smear the uncertainty that occurs at each step, warping both the communities and the interpretation of the dynamics.

The problems listed above can be mitigated using Temporal Communities through Trajectory Clustering (TCTC), which is designed to estimate temporal communities directly from node-collected data. TCTC bypasses the edge inference step and, instead, performs community detection in a single step directly from the time series. The solution here resembles ideas from the trajectory clustering literature (see Zheng, 2015 for a review), where their goal is to group trajectories in space and time.

There are numerous benefits to TCTC compared to existing methods. It can be used to discover the temporal and spatial properties of node-collected data. There are concrete hyperparameters that can be meaningfully tuned to identify settings for a contrast-of-interest. TCTC can also account for sudden spikes in noise that otherwise inherently bias methods that estimate network topology via a noisy edge-inference step. Finally, it identifies fluctuating temporal communities with a high temporal resolution, revealing new dynamic properties.

The article proceeds in the following manner: first, we introduce TCTC and demonstrate its utility through simulations, the recovery of expected time-averaged properties of task and resting-state neuroimaging data. After that, we show that the temporal information within TCTC contains information that relates to single-trial behaviour, revealing new information about ongoing temporal network configurations.

## Methods

### Description of TCTC

The TCTC algorithm identifies trajectories in a time series of nodes in a single step (see Figure 1A for an illustration of how this approach relates to other approaches). If nodes are part of a trajectory, they get assigned to a community. For nodes to be part of the same trajectory, they must comply with four rules: distance, size, time and tolerance (Figure 1B; Table 1). Briefly (see below for more detailed definitions), the distance rule states that all nodes in a trajectory must be within *ϵ* of each other. The size rule ensures the size of a trajectory contains at least *σ* nodes. The time rule ensures that the length of a trajectory must be *τ* or more time-points long. The tolerance rule states how many consecutive time-points can violate the previous three rules can while still allowing a trajectory to persist. Together, these hyperparameters define the minimum requirements for a community to exist within the data. These can be fit to find optimal hyperparameters by splitting the dataset into a training/test datasets (see N-back task below).

**Table 1:**
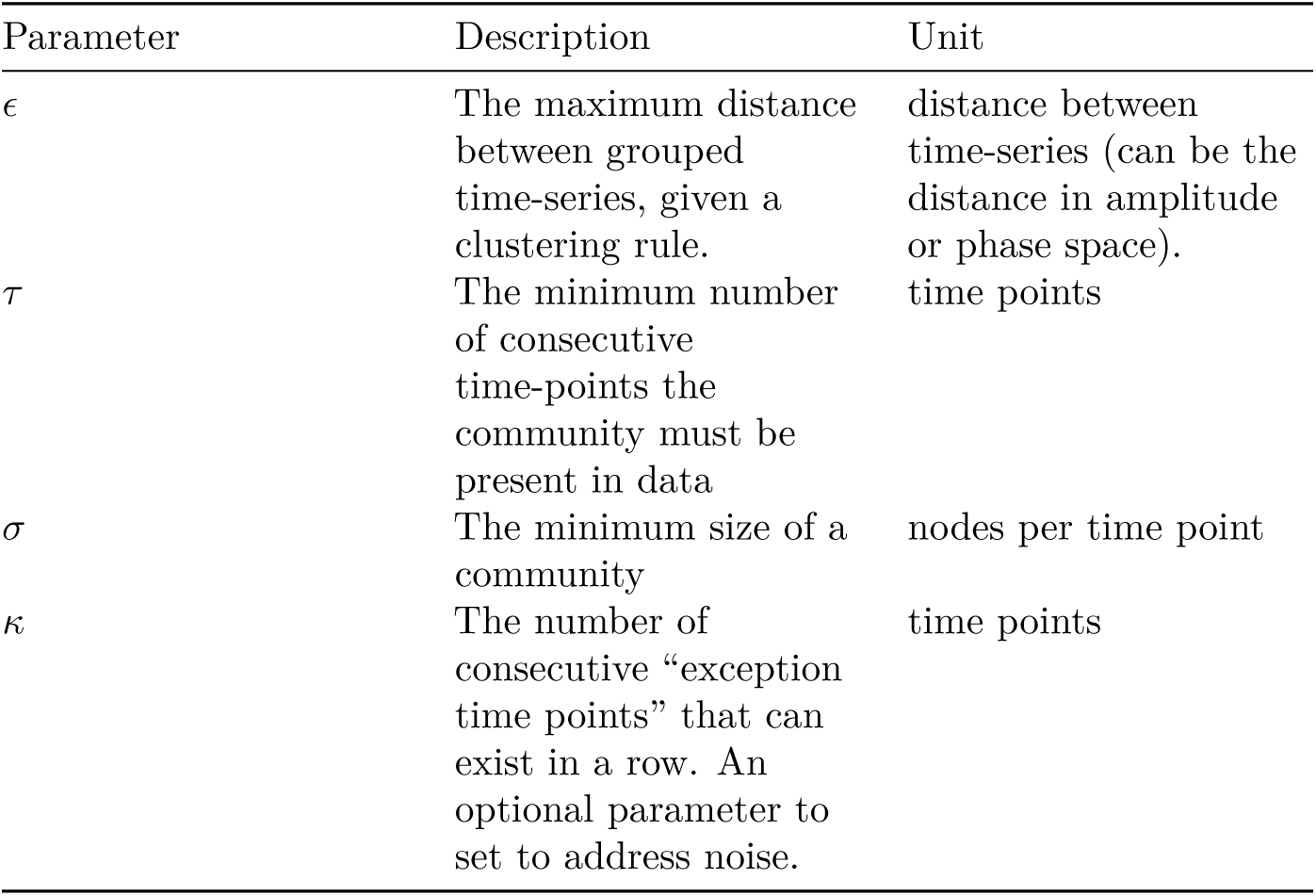
Description of hyperparameters involved in TCTC.

| Parameter | Description | Unit |
| --- | --- | --- |
| $\epsilon$ | The maximum distance between grouped time-series, given a clustering rule. | distance between time-series (can be the distance in amplitude or phase space). |
| $\tau$ | The minimum number of consecutive time-points the community must be present in data | time points |
| $\sigma$ | The minimum size of a community | nodes per time point |
| $\kappa$ | The number of consecutive “exception time points” that can exist in a row. An optional parameter to set to address noise. | time points |

**Figure 1:**
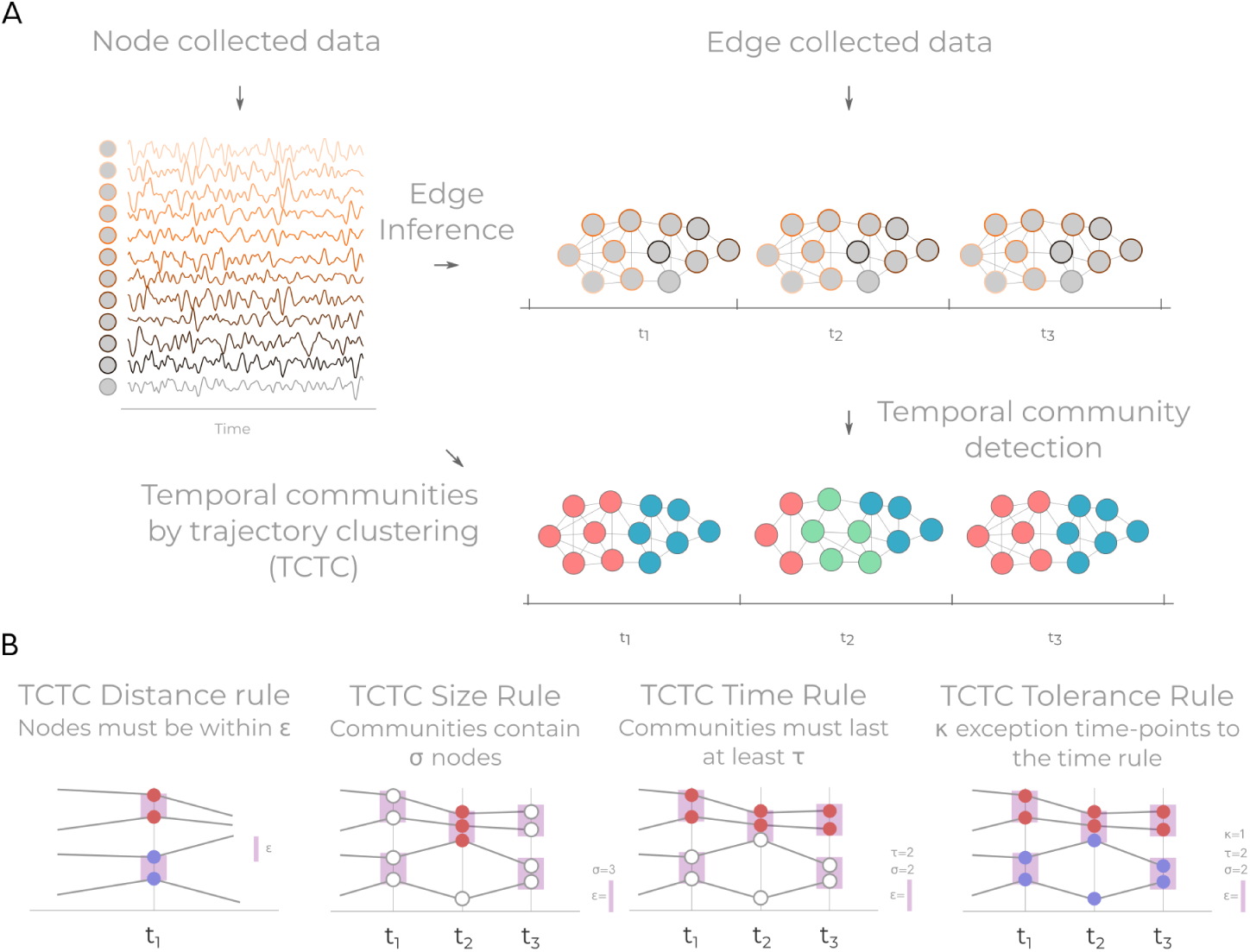
An illustration of TCTC. **A** How TCTC relates to other methods. Flow chart showing how temporal communities are derived contrasting TCTC from 2-step procedures on node collected data. Temporal communities derived on edge collected data is one step, node collected data in two steps via edge inference, and node collected data with TCTC in one step. **B** The four rules that define trajectories in TCTC. A trajectory must satisfy all four rules. Each rule shows four time-series with one or three discrete time-points. Purple rectangles show instances where the distance rule is successfully applied. White nodes indicate when no there was no trajectory identified.

More specifically, let *X* is a discrete time-series consisting of *N* nodes and *T* time points. *X* can express the amplitude or instantaneous phase of the nodes. The goal is to create communities for each time point. In TCTC, communities are identified between groups nodes if there the nodes are part of a trajectory. TCTC has four rules, each with their hyperparameter: a distance rule, a duration rule, a size rule, and a tolerance rule (Table 1). Here we use the notation 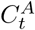 to notate “community A at time-point t” which consists of a set containing node indices that signify the nodes belong to that community.

TCTC is a multi-label community detection algorithm (Figure S2). This property entails that a node can belong to multiple communities at a single time-point and that communities can overlap, which is not common in many community detection algorithms used in neuroimaging contexts. This property can be advantageous if a node is accumulating information from multiple communities because it can become a member of each community with TCTC instead of forcing it to belong to a single community (or merging all communities into one).

### Distance rule

The distance rule specifies how far the different time series (in amplitude or phase space) are from each other. Given some distance function, *D*(*X*), the hyperparameter *ϵ* specifies the maximal allowed distance that time points can be from each other in order to be part of the same trajectory. When *ϵ* is small, only time series with very similar values with getting grouped in the same trajectory. As *ϵ* increases, more divergent time series will get clustered together. Explicitly, the rule is:

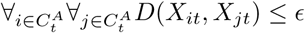

Where ∀*i* indicates “for all *i*”. This rule entails that the maximal distance between any node in a trajectory is *ϵ*. This formulation creates an analytic uncertainty of all the nodes within a community. For the distance function, we use *D*_1_ distance whenever *X* consists of amplitudes and, when *X* contains phase information the distance function is: *D* = |*X*_*jt*_ − *X*_*jt*_ | mod 2*π* (i.e. remainder of *D*_1_ after dividing by 2 *π*). The above formulation is similar to “flock clustering” from the trajectory clustering literature (Zheng, 2015).

### Duration rule

The duration parameter (*τ*) specifies the minimum length of the trajectory. This entails that the nodes that are a member of 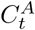 are also a member of 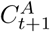 and all subsequent time-point up until 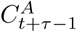. That is to say the community obeying the duration rule must follow:

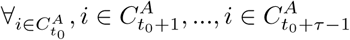

Where *t*_0_ signifies the first time-point in a trajectory.

### Trajectory size

The size parameter (*σ*) specifies the minimum number of nodes that are part of the trajectory. This parameter entails that there must be at least *σ* nodes belong in 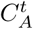. Explicitly, a community must follow the following size rule:

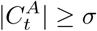

Where ‖ indicates the number of elements within the set.

### Tolerance rule

The tolerance rule specifies how many consecutive exceptions are allowed where the distance rule or size rule fails. The idea here is that, if a brief spike in noise affects one or more of the time series, this will interrupt the trajectory. If *τ* = 3 and *κ* = 1 then it is possible for there to be two instances where the tolerance rule can be applied (at *t*_0_ + 1 and *t*_0_ + 3). This results in all members of *C*^*A*^ being present at *t*_0_, *t*_0_ +2, *t*_0_ +4. This parameter is not necessary for TCTC to function. It is an additional parameter that is added to help mitigate problems with noise. Thus TCTC requires three parameters and has one optional parameter.

### Simulations

For the simulations, two sinusoids were generated:

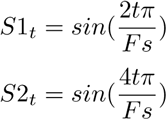

Where Fs was set to 100. Three time-series were generated by sampling from one of the sinusoids and adding Gaussian noise:

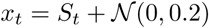

The length of each time series was 200. One time series only sampled from S1, the second time series only sampled from S2, and the third time-series switched between the two. The switches occurred at t=33, 67, 100, 133, 167. The ground truth assigned to each community at *t* was the sinusoid that the time series was following at *t*.

For simulation 2, one time-series was assigned a pulsating noise spike every 20 time-points (starting at time point 5) and a second time-series was assigned a pulsating noise spike every 35 time-points starting at time point 10. The amplitude of the noise spike was to add 2 * Beta(5, 1) (where Beta entails sampling from a beta distribution) to the time point generated value. The pulse lasted for two time-points.

Three different methods were tested. TCTC (distance rule applied to amplitude space), TCTC (distance rule applied to phase space) and a two-step method which involved a sliding window (boxcar taper) with a multi-layer Louvain (SW+MLL) (Mucha et al., 2010). All parameters for all three methods were fit with a grid search optimisation. A “training” and “test” simulation was created where each method could identify the best fitting parameters on the training simulation. The training was evaluated by the sum of the average normalised mutual information score at each time point. The search space the methods were: For TCTC(amplitude space) *τ* : 2-50 in steps of 2, *ϵ*: 0.1-1.5 in steps of 0.1, *σ*: 2-3, *κ*: 0-2. For TCTC(phase space) *τ* : 2-50 in steps of 2, 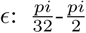 in steps of 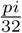, *σ*: 2-3, *κ*: 0-2. SW+MLL window size: 15-55 in steps of 10. The multi-layer parameters *γ* and *ω* were both between 0-1.5 in steps of 0.25.

### fMRI data

#### Dataset 1: Midnight scanning club data

For the resting-state analysis, there were ten subjects with ten resting-state sessions (818 time-points) from the Midnight Scanning Club (MSC) dataset (Gordon et al., 2017). One subject (MSC08) was deleted as is known to be noisy. The preprocessed data, as outlined in ref Gordon et al. (2017), available on OpenNeuro, was used. The only exception was that 200 parcels were created from the Schaefer atlas (Schaefer et al., 2018) and the Yeo 7-community static network parcellation (Yeo et al., 2011). Static functional connectivity was calculated with a Pearson correlation across each pairwise combination. For TCTC, the time series were first standardised to have a mean of 0 and a standard deviation of 1.

#### Dataset 2: HCP N-back task

One hundred subjects from the Human Connectome Project N-back task while recording fMRI (100 unrelated subject release, TR=0.72, minimal preprocessed data used) (Glasser et al., 2013; Van Essen et al., 2012). We used the LR encoding dataset throughout the paper except in the verification of the Bayesian models where we used the RL encoding. The data was split into training and test datasets, each containing 50 subjects.

The same 200 ROIs and seven static network partition that was used in the resting-state analysis were used here too. We regressed out 12 movement parameters, framewise displacement and global mean. Scrubbing was used to remove any time points that had a framewise displacement (FWD) value greater than 0.5. Missing data were simulated with a cubic spline to create a continuous time series. A subject would have been removed if more than 20% of a subjects data were simulated. No subjects met this criterion. The data was band-passed between 0.01 and 0.12 Hz, and the data was converted to instantaneous phase. Each subject performed eight blocks (four 0-back and four 2-back). These additional preprocessing was done using nilearn (Abraham et al., 2014) and Teneto (Thompson, 2019).

#### TCTC parameters

For the MSC resting-state data, preset parameters were chosen. These were: *τ* : 5, *ϵ* 0.5 (amplitude space), *σ*: 5, *κ*: 1. One of the reasons for choosing preset here is to demonstrate how these parameters are interpretable. Here we have stated that to be a community they must: last for five time-points, consist of at least five nodes, and those five nodes must always be within half a standard deviation of each other. Finally, there may be a single time-point where the previous conditions are not met. How these parameters shape the community detection algorithm is straight forward given the definitions. To demonstrate the sensitivity of changing these parameters, the effect of modifying each of the parameters is in Figure S3.

For the N-back task data, the objective function was defined to maximise the Hamming distance between binary trajectory clustering matrices. Each block was 33.12+*τ* seconds long (The block lasted from the first stimulus onset, 35 frames after stimulus onset, plus an additional ten frames to account for the sluggishness of the signal). When contrasting between sessions, the first ten time points were removed to avoid training on any spillover from the previous block. We arrived at these hyperparameters by optimising 50 subjects in the training dataset. Grid search optimisation search for the best performing hyperparameter combinations. The grid searched was 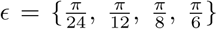, *σ* = {3-10}, *τ* = {3-10}, *κ* = {1,2,3}.

The goal of the optimisation was to minimise the following equation:

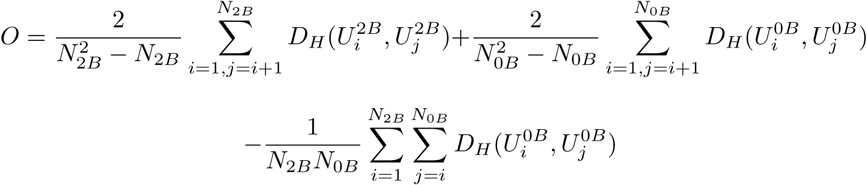

Where U is the upper-triangular of each temporal snapshot of the binary trajectory matrix (dimensions: node, node, time) where 1 signifies a trajectory is present. *D*_*H*_ is the hamming distance. 0B and 2B indicate 0-back or 2-black conditions and *i* and *j* are the condition index of U. *N*_2*B*_ is the number of blocks of the 2-back condition (here four). To minimise *O*, this entails that average difference between “within 2-back” and “within 0-back” blocks should have a small hamming distance and the average difference “between 2-back and 0-back” blocks should be as large as possible. The results of the optimisation are in Figure S6. From inspecting these results we chose *τ* : 5, 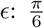, *σ*: 6, *κ*: 2 as there was a trough optimisation space here.

Note, we found communities based on the amplitude of the nodes on the MSC dataset. For the N-back, we identified communities based on the phase of the time series. We have done this to illustrate the two different possibilities for TCTC. Our preference is to use phase space, especially for task data, as any mean non-stationarity occurring in the data will affect the amplitude space communities more than phase space.

#### Community detection metrics

To ease the interpretation of our new method, we define a set of additional metrics to quantify the time-varying communities TCTC (see Figure S2 for visualisation of both metrics).

#### Pairwise trajectory ratio (PTR)

To summarise the amount of interaction between the identified communities through TCTC and the static network template, we derived the *pairwise trajectory ratio*. For each node pairing, we count the percentage of time-points that two nodes appeared in at least one community together.

#### Static community co-occurrence (SCC)

To summarise the amount of interaction between the identified communities through TCTC and the static community template, we derived the *static community co-occurrence*. For each static community pairing, we counted the percentage of nodes in all TCTC communities that intersect with the static functional network template from all possible nodes. Namely:

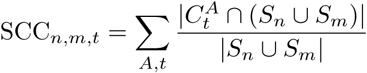

Where *S*_*n*_ and *S*_*m*_ are sets containing nodes indices for the static community partition for static network indices *n* and *m*. This measure is thus reflective of the overall intersection of the temporal communities, between the two static communities. When the term “within-SCC” it refers to the SCC_*n,n*_ and the “between-SCC” is the average SCC_*n,m,t*_ over all *m* when *n* ≠ *m*.

#### Statistics

Network-based statistic (NBS) was used (Zalesky, Fornito, and Bullmore, 2010) to determine significance between the two N-back blocks in the test dataset. NBS aims to find clusters of edges that significantly differ between conditions. Permutations of each block (4 per condition, per subject) entailed shuffling the condition-membership to create the permuted distribution. The PTR from each block were averaged over time entailing that any difference found is a time-averaged difference between the conditions. NBS ran for 1,000 permutations. We set the cluster threshold to 2 and significance was considered if p<0.001. This test derives a set of edges where the frequency of trajectories over time between different nodes were significantly different between the communities.

We used a hierarchical Bayesian model to quantify the difference between single-trial behaviour (reaction times, accuracy, and response). From the 28 SCCs, 5 PCA components (accounting for 75.5% of the variance) were derived expressing temporal network community configurations at each time-point during the block. The sampling of fMRI volumes does not correspond to stimulus onset. To account for this and to make sure we do not merely fit the stimulus jitter, a weighted average of the two encompassing PC time-points was used to align all trials to the same temporal offset. Each statistical model was run for the stimulus onset-locked PCs and up to 10 seconds afterwards. As different individuals have different reaction times for the different blocks (as each block had different stimuli types), each block had its own intercept modelled separately.

The statistical model that models single trial reaction times of correct trials from the community snapshots was specified as:

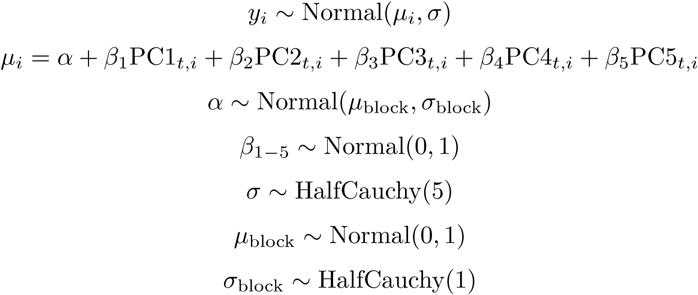

Where *y*_*i*_ was the reaction times. A Box-Cox transform was applied to the reaction times in order to transform them towards a Gaussian distribution (*λ* = 0.063, found using Scipy [V1.2.1, (Jones, Oliphant, Peterson, and Others, 2001)] Box-Cox function that finds the *λ* value that maximizes log-likelihood). The reaction times and PC components were standardised, so the *β* values are on comparable scales. All priors are weakly informative priors. This hierarchically models an intercept (*α*) for each of the blocks. MCMC was performed using pymc3 (Salvatier, Wiecki, and Fonnesbeck, 2016). Ten thousand samples were drawn from 3 separate chains (1,000 tuning samples for each) using the no-U-turns sampler (NUTS). The model ran for 11 different values of t (t=0, to t=10). For modelling both accuracy and response, a similar model was applied as above, except with small modifications to make the model logistic instead of linear to account for *y* being binary values where 0 was an incorrect or miss trial and 1 was a correct trial:

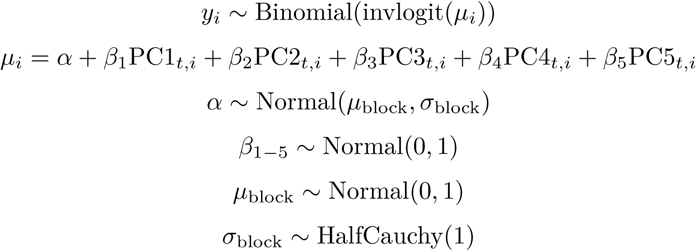

Evaluation checks of MCMC were done through manual inspection and checking that the Gelman-Rubin statistic was close to 1. The model selection used the leave-one-out information (LOO) criteria.

##### Verification of results on unseen data

An unseen dataset (RL encoding, see above) verified the best LOO model for each behaviour. Verification of models involved sampling from the posterior distribution and comparing to the verification dataset. Ten thousand samples of the posterior were drawn. For the linear model (reaction time), we calculated posterior predictive p-values between the simulated value and the original data for the mean and IQR. With the verification dataset, a linear model was fitted between the simulated data and unseen reaction times sampling all priors and distributions for similar to the linear model outlined above (where the model was: VerificationRTs ∼*α* + *β*\* PosteriorRTs. All priors were weakly informative). Both the independent and dependent variables were standardised. For logistic models, we first calculated a predictive threshold after viewing the ROC curves and selecting the maximum point that corresponded to: (1-false positives)+(true positives). We applied this threshold to the verification dataset, and the weighted F1 score was used to evaluate the predictive accuracy of the models.

## Code and Data availability

Code for TCTC is implemented in Teneto (https://github.com/wiheto/teneto) from v0.4.4 and onwards. Data for N-back task can be found on the Human Connectome Project homepage (https://humanconnectome.org/), and MSC dataset is available on Open Neuro (https://openneuro.org, ds000224).

## Results

### Validating TCTC with simulations

We create two simple simulations to illustrate TCTC’s ability to detect communities through time. We contrasted TCTC (using both phase and amplitude space) with the sliding-window method to derive time-varying connectivity and multi-layer Louvain (Mucha et al. (2010)) to derive the community detection (hereafter SW+MLL).

In each simulation, three time-series were generated, representing the activity in three nodes (Figure2AE). In simulation 2, there were moments where uncorrelated spiking noise artefacts occurred for the green and blue time series. The ground-truth community partition was based on when the underlying sinusoids that time series were sampled from overlapped (Figure 2BF).

**Figure 2:**
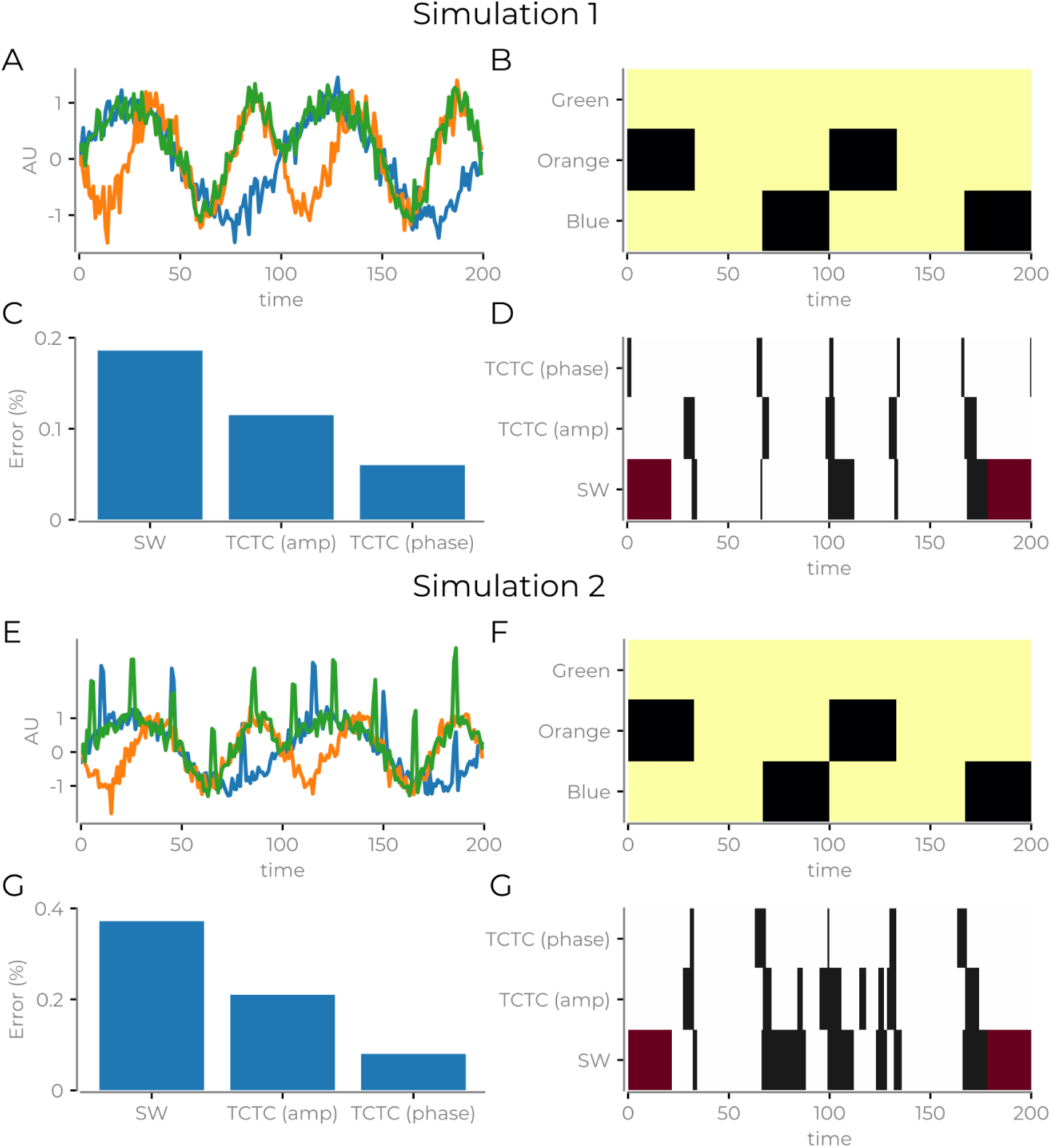
Results from simulations. **A** The time series generated in simulation one. **B** The “ground truth” for the different simulations. Different colours indicate different communities. **C** Percent of time-points where the method failed to identify the ground truth when applying the best fitting parameters from each method to unseen test data. For the SW + Louvain method, there were time-points where it was unable to calculate any estimate and excluded from the error calculation. **D** Illustration showing which time-points were errors (black), correct (white) and could not be calculated (red). **E** Time series generated for simulation 2. Aside from random moments of spiking noise inserted into two of the time series, all other parameters from simulation 1 were preserved. **F** Same as B, but for time series in simulation 2. **G** Same as C, but for the time series in simulation 2. (H) Same as D, but for the time series in simulation 2.

All methods had their hyperparameters fitted in a grid-search procedure (see Methods). After training, we tested the best fitting parameters against unseen test data. TCTC (phase) performed the best in both simulations (Figure 2), followed by TCTC (amplitude) and then the SW + Louvain method (despite being unable to quantify the entire time series) (Figure 1CG). Figure 1D and 1H show when the different methods failed to recover the ground truth, we see that TCTC (phase) had brief periods of errors. Whereas SW + Louvain suffered from more prolonged periods of error, especially when there was additional noise in the simulations. Even in these straightforward simulations, they still show the critical properties of temporal community splitting and merging (Granell, Darst, Arenas, Fortunato, and Sergio (2015)). In sum, TCTC can outperform SW + MLL.

### Validating TCTC on fMRI data

Here, we demonstrate the validity of TCTC when applied to fMRI data. In order to validate the approach, we first consider whether the time-averaged communities from TCTC reveal static connectivity properties commonly found during rest with fMRI (Fox et al., 2005; Fransson, 2005). If this is the case, then TCTC is identifying properties that, while they may be fluctuating through time, when pooled together recreate the expected static relationship. We illustrate this in both a resting-state fMRI and task fMRI datasets.

Using the MSC resting-state dataset, we found a clear relationship between pairwise trajectory ratio (PTR) and the static functional connectivity (Figure 2A). Further, we averaged the PTR for each static network combination (Yeo et al. (2011)), and find high similarity between the average PTR and average functional connectivity between the static communities (Figure 2BC). Here we see that TCTC identifies properties similar to resting-state networks. This is important to establish because, in absence of a ground truth, recovering known time-averaged properties is an important validating step. Furthermore, to verify that TCTC is indeed finding session-specific properties, we compare the PTR with the functional connectivity from (i) the same subject/sessions (median *ρ*: 0.54), (ii) other sessions from the same subject (median *ρ*: 0.42), and (iii) other subjects/sessions (median *ρ*: 0.29) (Figure 2D). Here we see higher correlations when TCTC is matched with the session’s corresponding functional connectivity and decreases as the expected relationship between the variables decrease. In sum, when averaging over time, TCTC recreates expected connectivity patterns at rest.

The preceding analysis validates that TCTC is sensitive to time-averaged signal within its session compared to other sessions. Next, we examined whether it is sensitive to expected time-average fMRI signals when performing a task. Here we use the data from the N-back task within the Human Connectome Project (HCP) dataset (Barch et al., 2013; Van Essen et al., 2012). Using similar logic to the previous verification, we consider whether there are time-averaged differences in communities that resemble expected differences in an N-back task (e.g. Barch et al. (2013); Finc et al. (2017)). The PTR differences between 2-back and 0-back in the test-dataset were used to identify which pairs of nodes that often ended up in the same community for a specific condition.

There were significant differences between the 0 and 2-back blocks on the test dataset (Figure 3AB, NBS statistics, p<0.001, cluster threshold: 2). The communities identified in the 2-back block relate more to attentional, visual and control areas, whereas those identified in the 0-back task relate to the default mode network’ connectivity with other communities (Figure 3CD). Considering the subset of PTR combinations that were more frequent in the 2-back block, we observed a sustained period of activation throughout the block where there are more nodes throughout the entire time series in the 2-back condition (Figure 3EF). The reverse trend exists for the PTR during the 0-back block. This result demonstrates that the block differences (Figure 3AB) not driven by a handful of time-points, but sustained throughout the blocks. However, this result does not entail that there are no fluctuations in the temporal communities during the block.

**Figure 3:**
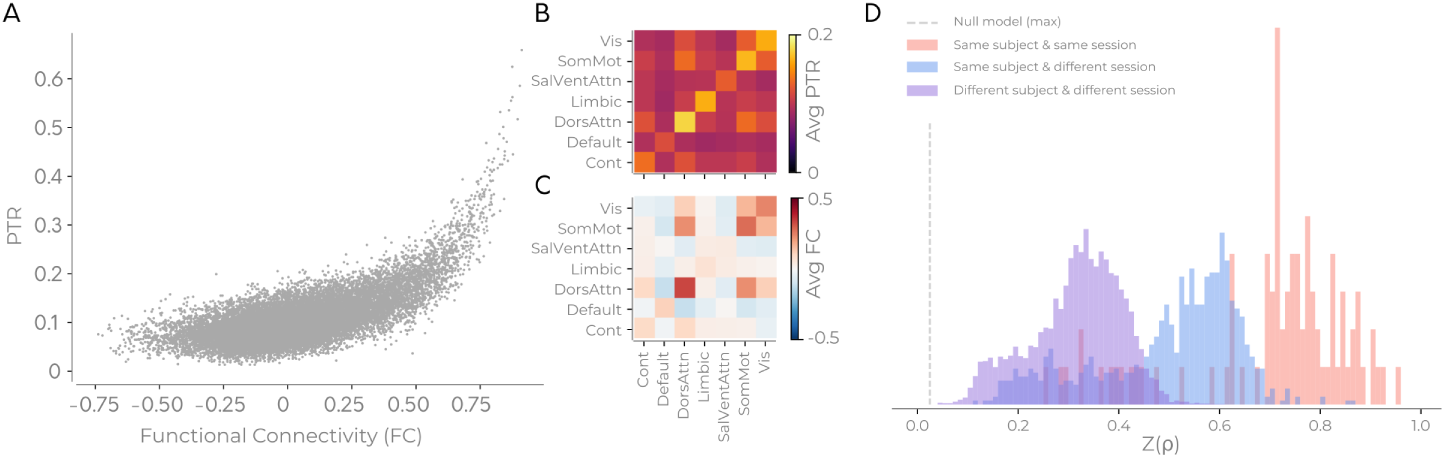
Verifying TCTC on the MSC dataset. **A** Relationship of each edge’s static functional connectivity and the ratio of time-points that two nodes get classed in the same community for an example subject/session. **B** Using a static brain network template, the average number of trajectories for each brain network combination from the example subject in *A*. **C** Same as *B* but for functional connectivity. **D** “Same session” shows the distribution of the correlation (Fisher transformed Spearman values) shown in *A* but for all subjects/sessions (same session). “Same subject” entails the distribution of pairwise trajectory ratio when correlated with the functional connectivity of different sessions from the same subject. “Different subjects/sessions” shows the distribution of trajectory numbers when correlated with the functional connectivity of different subjects/sessions. *D* shows all possible permutations for the different groups. Dashed lines in *D* are the maximal value when permuting the functional connectivity edges that get paired with the PTR 100 times for each subject.

### TCTC identifies fluctuating communities that have single-trial properties

The previous section showed that the average TCTC information contains relevant signal both in regards to task differences and subject properties during rest. However, the temporal specificity of TCTC entails it may be useful to identify event-related effects with greater temporal precision. This is the novelty of TCTC. In this section, we demonstrate that the temporal community fluctuations correlate with single-trial behavioural properties, and thus showing that they are not merely noise.

In order to illustrate the properties of the temporal information, we derive SCC values relative to stimulus onset for each 2-back trial and correlate this with single-trial response times and accuracy. We interpret t<4 to reflect prestimulus activity due to the sluggishness of the BOLD response.

We first reduced the 28 network configurations to five PC components (Figure 5AB, accounting for 75.5% variance). The five PC components each represent different temporal network configurations expressed along the static template dimensions (Figure 5AB). PC1 shows a general increase in all communities. PC2 is a community configuration containing more nodes from the visual network (both with other visual network nodes and with attention and sensorimotor networks). PC3 is marked by an increase in the limbic network increasing. PC4 shows an increase in communities containing nodes from the dorsal attention network. PC5 shows an increase in communities between cognitive control, dorsal attention and limbic networks. There are also fewer nodes from the default mode network in the communities. Together, these five components show a diverse number of community assignments.

**Figure 4:**
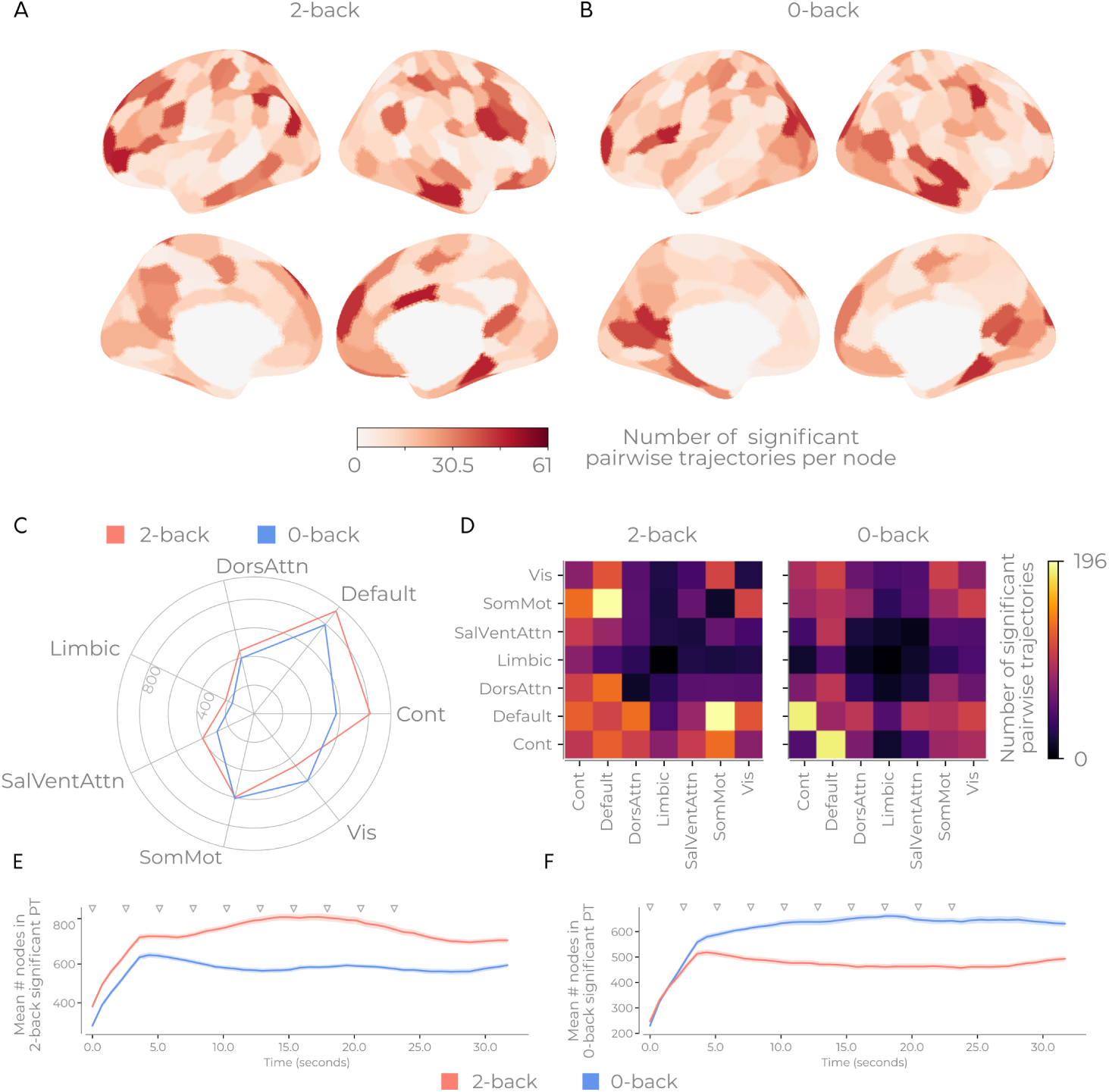
Verifying TCTC on HCP N-back task. **A** Differences between the number of trajectories assigned to a node differs between the 0 and 2-back tasks. **B** Placing the total trajectory differences between the two tasks onto the static information for the significant trajectories in the 2-back condition. **C** Same as *B* but tor the 0-back condition. **D** Displaying when in time, the two sets of communities in **B** are present for both tasks. **E F**

**Figure 5:**
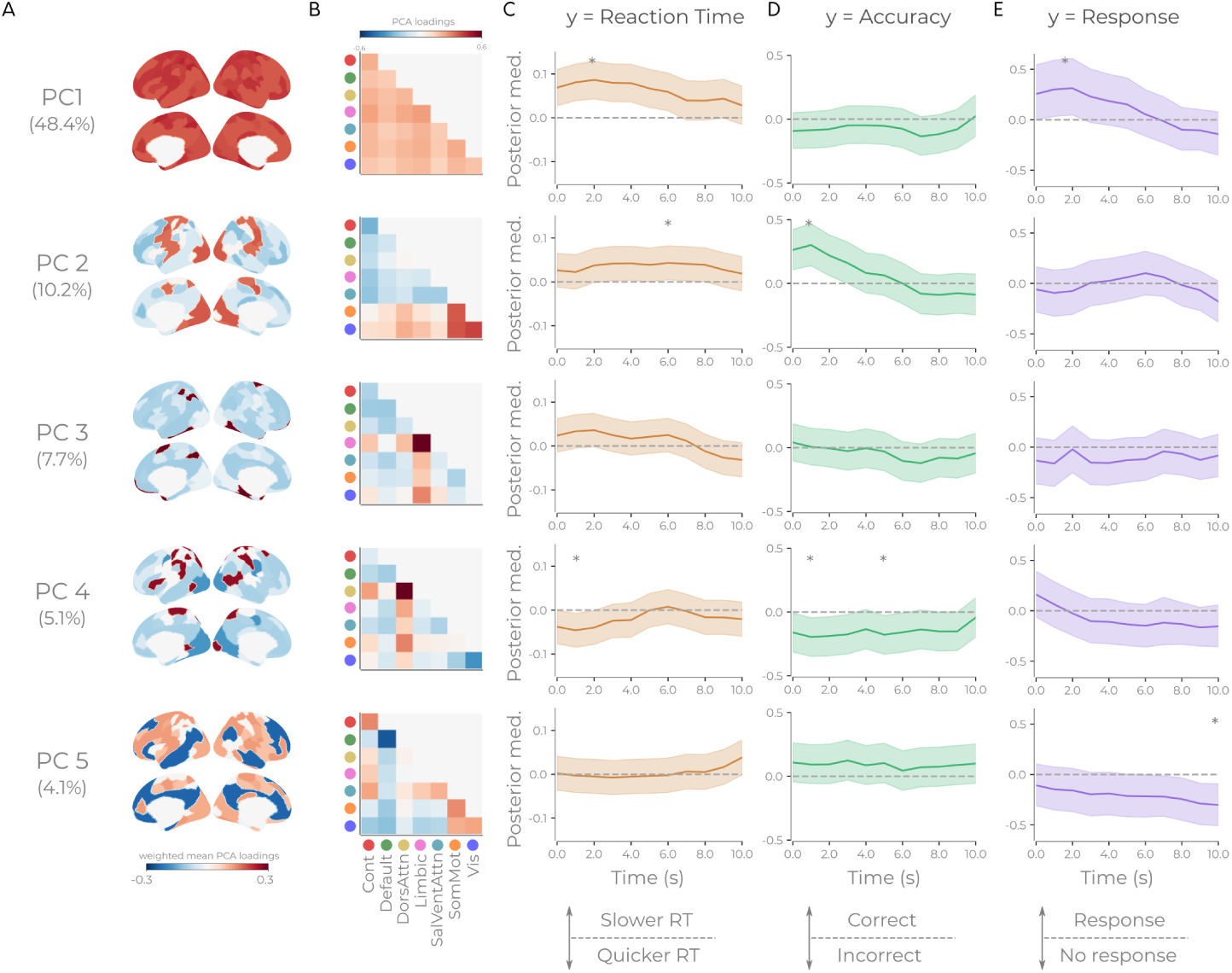
Snapshots of temporal community configurations correlate with single-trial behaviour. **A** Weighted average PCA loadings where the “within” static network loading is weighted as much as all the “between” static networks together. **B** PCA loadings when reducing the network combination for each trial to 5 PCA components for the SCC values at around each trial onset. **C** Linear model where the SCC PC components model the reaction times. Lines show posterior medians of each PC for Bayesian models fitted for each time-point relative to stimulus onset. The shaded region shows the 90% credible interval around the mean. Asterisks depict max/min median posterior value through time where the entire 90% credible interval peaks either above/below zero. Information of the posterior distribution at each astrisk is detailed in Table S4. **D** Same as *C* but for a logistic model modelling accuracy. **E** Same as *C* but a logistic model modelling whether there was a response or not.

The results reveal that the temporal network configurations are associated with behaviour differently depending on when they occur (Figure 5CDE). We highlight the peaks/troughs in the posterior distributions where the credible intervals are 90% above/below zero. An increase in PC1, (i.e. global integration between the static brain networks), before stimulus onset, is associated both with slower reaction times but also ensuring a response happens (Figure 5CE).

PC2, which involves the visual network being in communities with other brain networks, gave more correct trials before stimulus onset but slower reaction times if it persisted. Quicker reaction times were associated with reduced PC4 before stimulus onset and also lead to more inaccuracies both before and after stimulus onset (Figure 5CD). This attentional component seems to be involved in the speed-accuracy trade-off. Finally, when PC5 is low (i.e. more default mode network nodes involved in communities), and was involved later in the trial, it entailed a response was absent. This result seems to be indicative of task-interfering processes, such as mind-wandering, occurring (Figure CE).

A detailed picture emerges in which different network interactions, at various times, can explain multiple behavioural measures. This result confirms that TCTC is sensitive to behaviourally-relevant temporal information within neuroimaging data. However, as each time-point has received its model, we have yet to show if multiple time-points together explain single-trial behaviour. We selected the features where a network combination peaks where 90% of the credible interval is above/below 0 into one model per behaviour. The LOO reveals that combined models were the best performing (see Table S1-S3). Posterior distributions for the combined models are shown in Figure 6A-C where their results illustrate how different components, at different time-points, account for that behaviour. In sum, the different TCTC configuration at multiple time-points together can explain single-trial behavioural properties.

**Figure 6:**
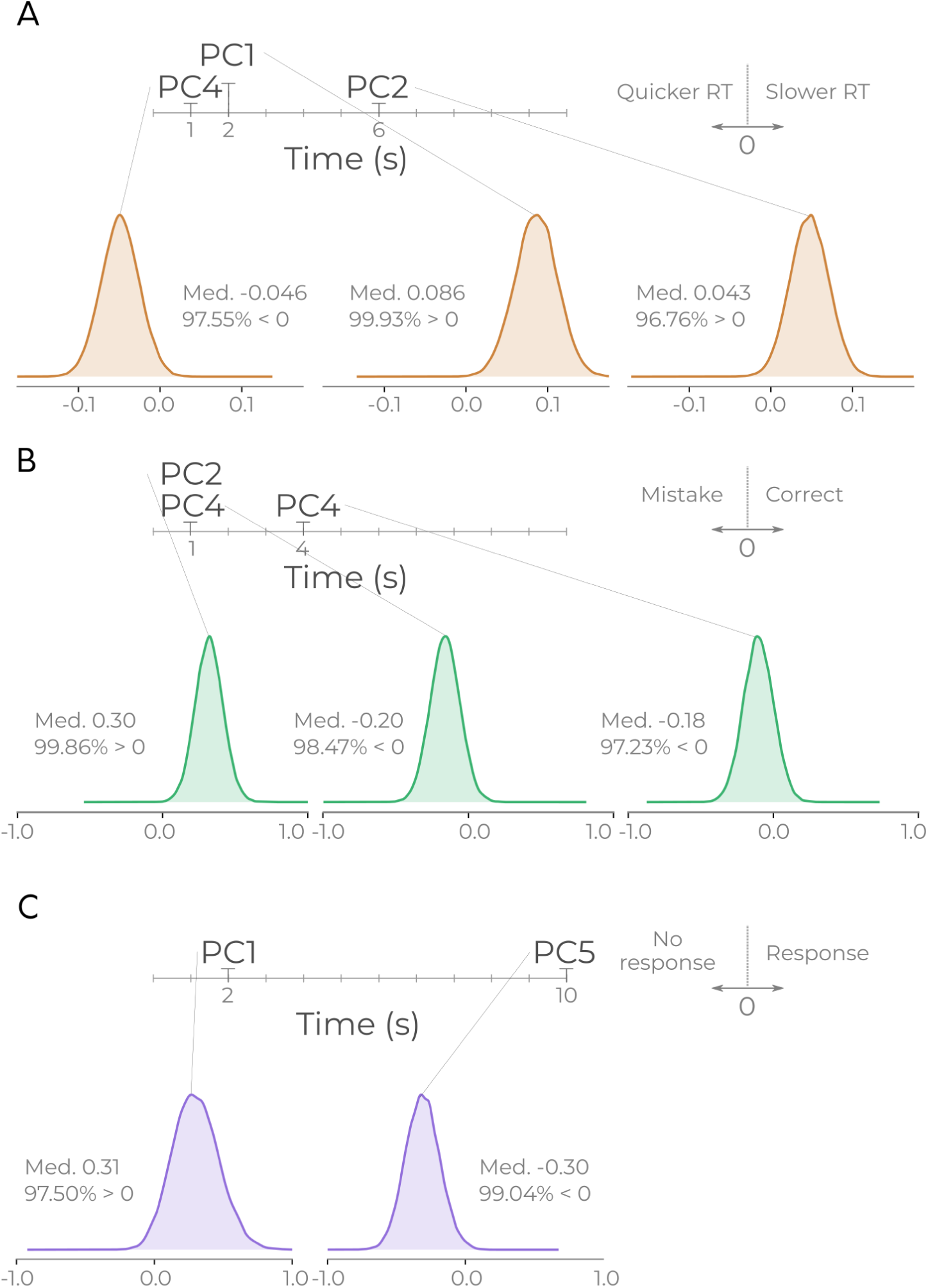
Posterior distributions of multi-time point models show multiple temporal network configurations correlate with single-trial behaviour. **A** The posterior distributions of three PC component selected for the combined reaction time model. Lines from each distribution point to which PC component and when in time. **B** Same as *a* but for three selected components for accuracy. **C** Same as *A* but for the two selected component for response model

#### Verification of combined models on unseen data

Finally, in order to verify the selected models for each behaviour, we sample from the posterior distribution and compare the sampled data to both the original data (posterior predictive checks) and an unseen verification dataset (prediction). For the linear model for reaction time, the mean and interquartile range (IQR) of the simulated data were compared with the original data (mean: p=0.38, IQR: p=0.21) indicating a good model fit which should generalise to new data. With the separate verification data, there was a correlation between the average posterior sampled data and the new verification data’s reaction times (median=0.27, 90% CI=[0.23, 0.31], 100% posterior above 0, Figure S4). This result shows that the model generalises to new data but also emphasises that the model is only capturing part of the variance of single-trial reaction times. In regards to the logistic models, we calculated the weighted F1 score of assigning a trial to an outcome. Due to the small number of errors and missed responses, we first identify a cut-off threshold for prediction by inspecting the ROC curve after sampling the posterior distribution. We then use this cut-off on when sampling from the posterior and comparing with the verification dataset. Both accuracy and response models had a high weighted F1 score (accuracy: original data: 0.81; verification data: 0.71; response: original data: 0.81; verification data: 0.75). In sum, all models show that their posterior distributions can model new unseen data.

## Discussion

Here we have introduced TCTC, a multi-label community detection algorithm designed for node collected data, and demonstrated the utility of the algorithm on two separate functional neuroimaging datasets. Critically, it can estimate community structure directly from time-series, without requiring additional estimates of network edges. We have shown: (i) TCTC outperforms 2-step methods in simple simulation (ii) that the hyperparameters from TCTC are interpretable; (iii) that expected time-averaged results are recoverable from time-averaged communities; and (iv) that the temporal information identified in the communities correlates with single-trial behaviour during a cognitive task. Thus, TCTC can probe how interacting sets of communities cooperate through time during a task.

We have demonstrated with TCTC that multiple network configurations are behaviourally relevant at several moments and that the same components can affect multiple behaviours. This result signifies that the ongoing community configurations are in flux, and these configurations are essential for efficient transfer of information across the brain. This model suggests a continual interplay of interactions across the traditionally static brain networks when performing a task. We have also demonstrated novel findings with TCTC regarding temporal communities and neuroimaging. Previously, researchers have quantified how average performance relates aggregate measures of temporal communities (e.g. Bassett et al. (2011); Shine, Koyejo, and Poldrack (2016); Saggar et al. (2018)) comparing rest to task data (Mattar, Cole, Thompson-Schill, and Bassett (2015)), all using two-step methods. Usually, behavioural correlates of network measures are on averaging the behaviour over trials (e.g. Saggar et al. (2018)). Here we have demonstrated that roles can change during a time series and the importance of temporal communities.

This single-trial time-varying finding contrasts with many analysis protocols that instead aim to identify brain regions, patterns, networks, or network configuration that are considered “on” or “more greatly activated” during a task condition. Instead, it suggests we should view cognitive processes in terms of information flow in the brain occurring between communities that merge and split based on the task at hand. If the right nodes interact and at the right time, the correct information flows around the brain and, in turn, will lead to greater accuracy and quicker reaction times. To perform a task, the dynamic coordination of multiple brain regions can affect performance, but only at the correct time, otherwise, they can be detrimental. Such temporal zones of useful connectivity configurations lead to new possible hypotheses regarding how large scale networks should be quantified and how they get attributed for different cognitive processes. Primarily, this perspective suggests pivoting the field away from identifying brain areas/networks/network configurations that are merely “on” or “off” more during a task, and towards identifying the spatiotemporal configurations of networks as they facilitate information flow in the brain. In sum, the results from TCTC opens up an avenue of research questions to explore quick changes in time-varying communities in neuroimaging contexts.

One possible concern regarding TCTC is its complexity. The hyperparameters themselves are simple. However, an algorithm with four hyperparameters is quite complex, considering other approaches have fewer (e.g. Mucha et al. (2010) has 2). This concern seems to make TCTC more complex. However, other methods require additional steps before applying the community detection algorithm, nullifying this criticism as the degrees of freedom and parameters in the 2 step approaches are considerably larger. For example, if choosing to use a sliding window method (degree of freedom), there is an additional implicit degree of freedom regarding the taper used for the window, and the window’s parameters (e.g. the simplest taper, a boxcar, will have a window length parameter). Thus, the sliding window method with temporal community detection using multi-layer communities from Mucha et al. (2010) has three hyperparameters and multiple additional methodological choices to make. Thus, TCTC is not considerably more complex than other methods.

Various modifications to TCTC are possible, which may be appropriate for different use-cases. At present, lagged relationships are not present. Modifying the distance measure to dynamic time warping would be a possible way to include such relationships. Other preprocessing steps exist in the trajectory clustering literature, such as an initial compression of the time series, which can speed up calculations. There are also multiple additional clustering algorithms from the trajectory clustering literature (convoys, swarms), which could be applied as the distance rule (Zheng, 2015). These alternative clustering methods for the distance rule can increase the speed of TCTC in larger networks. Another possible extension is to add the time-series of confounds into the community detection algorithm (e.g. global signal). If a community also contains these confounds, discarded that community.

One final noteworthy property of TCTC is that the multi-label communities can be converted to create binary time-varying connectivity matrices. Such a transform could is possible with no additional loss of information (i.e. from the community to connectivity representations). This property opens up additional possibilities for time-varying connectivity through trajectory clustering (TVCTC) and performs analyses beyond community detection.

## Acknowledgments

WHT acknowledges support from the Knut och Alice Wallenbergs Stiftelse (SE) (grant no. 2016.0473, http://kaw.wallenberg.org). This Midnight Scanning Club dataset was obtained from the OpenNeuro database. Its accession number is ds000235. Human connectome data was provided by the Human Connectome Project, WU-Minn Consortium (Principal Investigators: David Van Essen and Kamil Ugurbil; 1U54MH091657) funded by the 16 NIH Institutes and Centers that support the NIH Blueprint for Neuroscience Research; and by the McDonnell Center for Systems Neuroscience at Washington University.”

## Reference

Abraham, A., Pedregosa, F., Eickenberg, M., Gervais, P., Mueller, A., Kossaifi, J., … Varoquaux, G. (2014). Machine learning for neuroimaging with scikit-learn. Frontiers in Neuroinformatics, 8 (February), 14. https://doi.org/10.3389/fninf.2014.00014

Barch, D. M., Burgess, G. C., Harms, M. P., Petersen, S. E., Schlaggar, B. L., Corbetta, M., … Van Essen, D. C. (2013). Function in the human connectome: Task-fMRI and individual differences in behavior. NeuroImage, 80, 169–189. https://doi.org/10.1016/j.neuroimage.2013.05.033

Bassett, D. S., Wymbs, N. F., Porter, M. a, Mucha, P. J., Carlson, J. M., and Grafton, S. T. (2011). Dynamic reconfiguration of human brain networks during learning. Proceedings of the National Academy of Sciences of the United States of America, 108 (18), 7641–6. https://doi.org/10.1073/pnas.1018985108

Bazzi, M., Porter, M. a., Williams, S., McDonald, M., Fenn, D. J., and Howison, S. D. (2016). Community detection in temporal multilayer networks, and its application to correlation networks. Multiscale Modeling & Simulation, 14 (1), 1–41. https://doi.org/10.1137/15M1009615

Finc, K., Bonna, K., Lewandowska, M., Wolak, T., Nikadon, J., Dreszer, J., … Kühn, S. (2017). Transition of the functional brain network related to increasing cognitive demands. Human Brain Mapping, 38 (7), 3659–3674. https://doi.org/10.1002/hbm.23621

Fortunato, S., and Hric, D. (2016). Community detection in networks: A user guide. Physics Reports, 659, 1–44. https://doi.org/10.1016/j.physrep.2016.09.002

Fox, M. D., Snyder, A. Z., Vincent, J. L., Corbetta, M., Van Essen, D. C., and Raichle, M. E. (2005). The human brain is intrinsically organized into dynamic, anticorrelated functional networks. Proceedings of the National Academy of Sciences of the United States of America, 102 (27), 9673–8. https://doi.org/10.1073/pnas.0504136102

Fransson, P. (2005). Spontaneous low-frequency BOLD signal fluctuations: An fMRI investigation of the resting-state default mode of brain function hypothesis. Human Brain Mapping, 26 (1), 15–29. https://doi.org/10.1002/hbm.20113

Glasser, M. F., Sotiropoulos, S. N., Wilson, J. A., Coalson, T. S., Fischl, B., Andersson, J. L., … Jenkinson, M. (2013). The minimal preprocessing pipelines for the Human Connectome Project. NeuroImage, 80, 105–24. https://doi.org/10.1016/j.neuroimage.2013.04.127

Gordon, E. M., Laumann, T. O., Gilmore, A. W., Petersen, S. E., Nelson, S. M., Dosenbach, N. U. F., … Greene, D. J. (2017). Precision Functional Mapping of Individual Human NeuroResource Precision Functional Mapping of Individual Human Brains. Neuron, 1–17. https://doi.org/10.1016/j.neuron.2017.07.011

Granell, C., Darst, R. K., Arenas, A., Fortunato, S., and Sergio, G. (2015). Benchmark model to assess community structure in evolving networks. Physical Review E, 92 (1), 012805.

Holme, P., and Saramäki, J. (2012). Temporal networks. Physics Reports, 519 (3), 97–125. https://doi.org/10.1016/j.physrep.2012.03.001

Hric, D., Darst, R. K., and Fortunato, S. (2014). Community detection in networks: Structural communities versus ground truth. Physical Review E - Statistical, Nonlinear, and Soft Matter Physics, 90 (6), 1–19. https://doi.org/10.1103/PhysRevE.90.062805

Jones, E., Oliphant, T., Peterson, P., and Others. (2001). SciPy: Open source scientific tools for Python.

Mattar, M. G., Cole, M. W., Thompson-Schill, S. L., and Bassett, D. S. (2015). A Functional Cartography of Cognitive Systems. PLOS Computational Biology, 11 (12), e1004533. https://doi.org/10.1371/journal.pcbi.1004533

Mucha, P. J., Richardson, T., Macon, K., Porter, M. a, and Onnela, J.-P. (2010). Community structure in time-dependent, multiscale, and multiplex networks. Science (New York, N.Y.), 328 (5980), 876–8. https://doi.org/10.1126/science.1184819

Newman, M. (2010). Networks. An introduction (p. 772).

Peixoto, T. P., and Rosvall, M. (2017). Modelling sequences and temporal networks with dynamic community structures. Nature Communications, 8 (1), 1–12. https://doi.org/10.1038/s41467-017-00148-9

Rosvall, M., and Bergstrom, C. T. (2010). Mapping change in large networks. PloS One, 5 (1), e8694. https://doi.org/10.1371/journal.pone.0008694

Saggar, M., Sporns, O., Gonzalez-castillo, J., Bandettini, P. A., Carlsson, G., Glover, G., and Reiss, A. L. (2018). Towards a new approach to reveal dynamical organization of the brain using topological data analysis Manish. Nature Communications, 9 (1399), 1–14. https://doi.org/10.1038/s41467-018-03664-4

Salvatier, J., Wiecki, T. V., and Fonnesbeck, C. (2016). Probabilistic programming in Python using PyMC3. PeerJ Computer Science, 2, e55. https://doi.org/10.7717/peerj-cs.55

Schaefer, A., Kong, R., Gordon, E. M., Laumann, T. O., Zuo, X.-N., Holmes, A., … Yeo, B. T. T. (2018). Local-Global Parcellation of the Human Cerebral Cortex From Intrinsic Functional Connectivity MRI. Cerebral Cortex, 28, 3095–3114. Retrieved from http://www.biorxiv.org/content/early/2017/06/06/135632

Shine, J. M., Koyejo, O., and Poldrack, R. A. (2016). Temporal metastates are associated with differential patterns of time-resolved connectivity, network topology, and attention. Proceedings of the National Academy of Sciences, 113 (35), 9888–9891. https://doi.org/10.1073/pnas.1604898113

Thompson, W. H. (2019). Teneto. https://doi.org/10.5281/zenodo.2535994

Van Essen, D. C., Ugurbil, K., Auerbach, E., Barch, D., Behrens, T. E. J., Bucholz, R., … Yacoub, E. (2012). The Human Connectome Project: a data acquisition perspective. NeuroImage, 62 (4), 2222–31. https://doi.org/10.1016/j.neuroimage.2012.02.018

Yeo, B., Krienen, F., Sepulcre, J., Sabuncu, M., Lashkar, D., Hollinshead, M., … Buckner, R. L. (2011). The organization of the human cerebral cortex estimated by intrinsic functional connectivity. Journal of Neurophysiology, 106, 1125–1165. https://doi.org/10.1152/jn.00338.2011.

Zalesky, A., Fornito, A., and Bullmore, E. T. (2010). Network-based statistic: identifying differences in brain networks. NeuroImage, 53 (4), 1197–207. https://doi.org/10.1016/j.neuroimage.2010.06.041

Zheng, Y. (2015). Trajectory Data Mining: An Overview. ACM Trans. On Intelligent Systems and Technology, 6 (3), 1–41. https://doi.org/10.1145/2743025

