## Supplementary Information for "The identification of temporal communities through trajectory clustering correlates with single-trial behavioural fluctuations in neuroimaging data"

### Supplementary Methods

#### Walkthrough of each TCTC parameter

Let us look at an illustration how each of the different hyperparameters work and how communities are inferred from the trajectories. If $X$ contains two time series, they get clustered together if they are within $\epsilon$ distance of each other given a distance function (Figure S1A), see methods for distance functions used). However, stationary time series will often cross each other incidentally, leading to brief moments where the two unrelated time series would be incorrectly grouped in the same community. Adding a minimum time requirement, $\tau$, (Figure S1B) assures that the communities persist for a minimum number of time points. If we now add a third time series to $X$, more community structures will be possible across the time series (Figure S1C). However, we may want communities to be of a minimum size; increasing the parameter $\sigma$ entails that a community consists of $\sigma$ number of nodes (Figure S1D). Finally, it is possible that time series can contain brief bursts of uncorrelated noise (Figure S1E). When increasing the final hyperparameter $\kappa$, it allows for some time-points to briefly violate the other rules (Figure S1E). In sum, these four different hyperparameters can set the spatial and temporal resolution of the identified communities. One of the benefits of this approach is these four different parameters each have a clear interpretation of how they affect the community detection.

#### Disclosure of additional statistical tests run

Prior to evaluating the Bayesian model for each second following stimulus onset in the hierarchical Bayesian model, two models were run ("reaction time" and "response" models (the latter being slightly misspecified as it also included incorrect trials in the "response present")). The priors were as outlined above for their respective models. The difference to this approach was threefold: (1) PC components were derived for only a portion of the block (1 per trial) instead of deriving the PC components first; (2) Analysis was performed only at 7.2 seconds after stimulus onset; (3) Only "reaction time" and "response" models were run. Regarding (1), this resulted with little difference regarding the PC components in regards to which networks had high loadings and overall variance explained. This pipeline lead to results which can be seen in figure 5. PC3 (for response) and PC4 (for RT) both had over 99% of their posterior distributions below 0. The motivation for changing the analysis was done after conceiving the approach performed in the main article (i.e. using the time dimension) and not due to the lack of success of this initial approach. The verification dataset was not touched with this analysis. Thus, only for the purpose of transparency of exploration do we disclose this.

Additionally, the entire analysis was run again, after correcting for error identified in the code and applying a simpler optimization procedure due to complexity concerns raised after the first version of the preprint of the article was released (the errors were the bandpass filter applied was different frequencies than specified in the text and some nodes assigned to incorrectly static community labels). After correction and substitution of the optimization, slightly different results were found (especially Figure 5). This has no significant change regarding the interpretation, significance of validity of TCTC. Aside from the change in optimization procedure, no changes in the procedure were made when the analysis was rerun.

#### Supplementary Figures

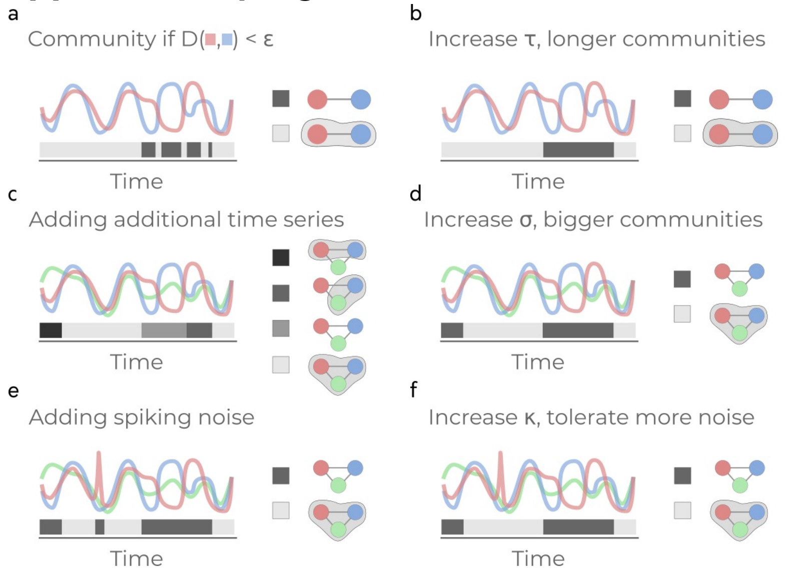


**Figure S1. Illustration of the different hyperparameters in TCTC and community inference from trajectories**. Each panel shows time series which are assumed to be node collected data from a network. The coloured bar along the bottom indicates the community assignment for that time point. The right side of each panel indicates the community structure where grouped nodes are joined together with shaded background **a** Two time series where $\tau$ is low are considered members of the same community if the distance between the two time series is less than $\epsilon$. **b** What happens to a when $\tau$ is increased. c Adding an additional time series to b and $\sigma = 2$. **d** The effect on c when increasing $\sigma$. **e** c has now been modified with a brief burst of noise. **f** The effect to the communities in e when increasing $\kappa$. This step ignores $\kappa$ number of time-points when the previous rules are not met.

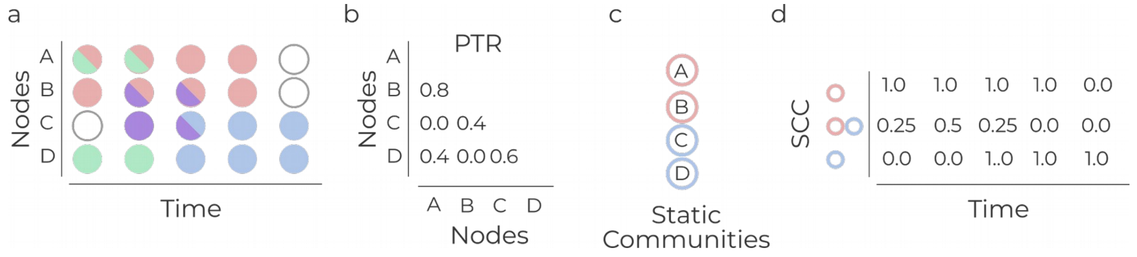


**Figure S2. Example of PTR and SCC to quantify the multi-label time-varying community assignments**. **a** an example of time-varying community assignment for five time-points. Time-points where a node has a white circle are when no community is assigned. Half circles indicate multiple communities assigned to the node. **b**. Pairwise trajectory ratio (PTR) for each of the node pairings in a. **c** A static community template for the nodes in a. **d** Given the static community template in c, the static community cooccurrence (SCC) is calculated. This is a value for each static community combination (both "within" a single static community and "between" pairwise combinations).

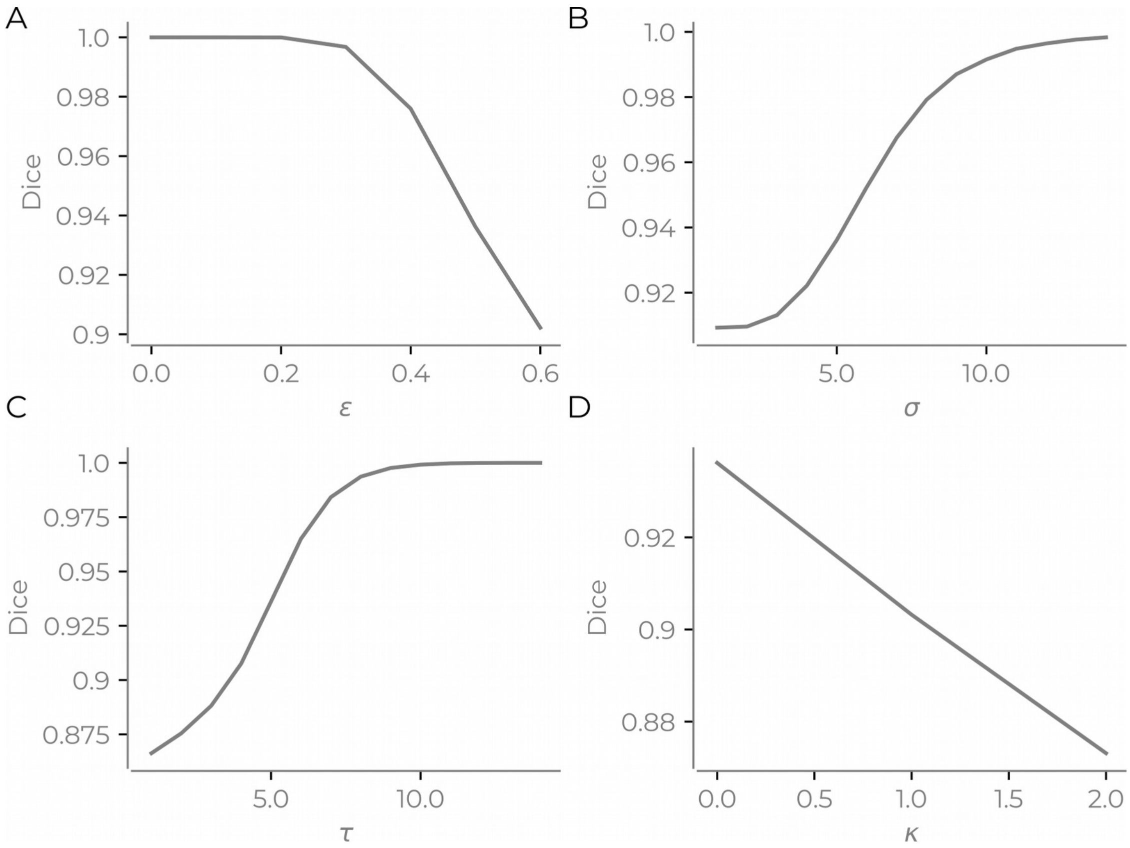


**Figure S3. Example of changing hyperparameters of TCTC for one MSC subject/session.** All variables are fixed at $\epsilon=0.5$, $\tau=5$, $\sigma=5$, $\kappa=0$ except for the variable that changes in each panel. **a** $\epsilon changes$, **b** $\sigma$ changes, **c** $\tau$ changes, **d** $\kappa$ changes. y-axis shows the average dice coefficient of each TCTC trajectories with the static community template.

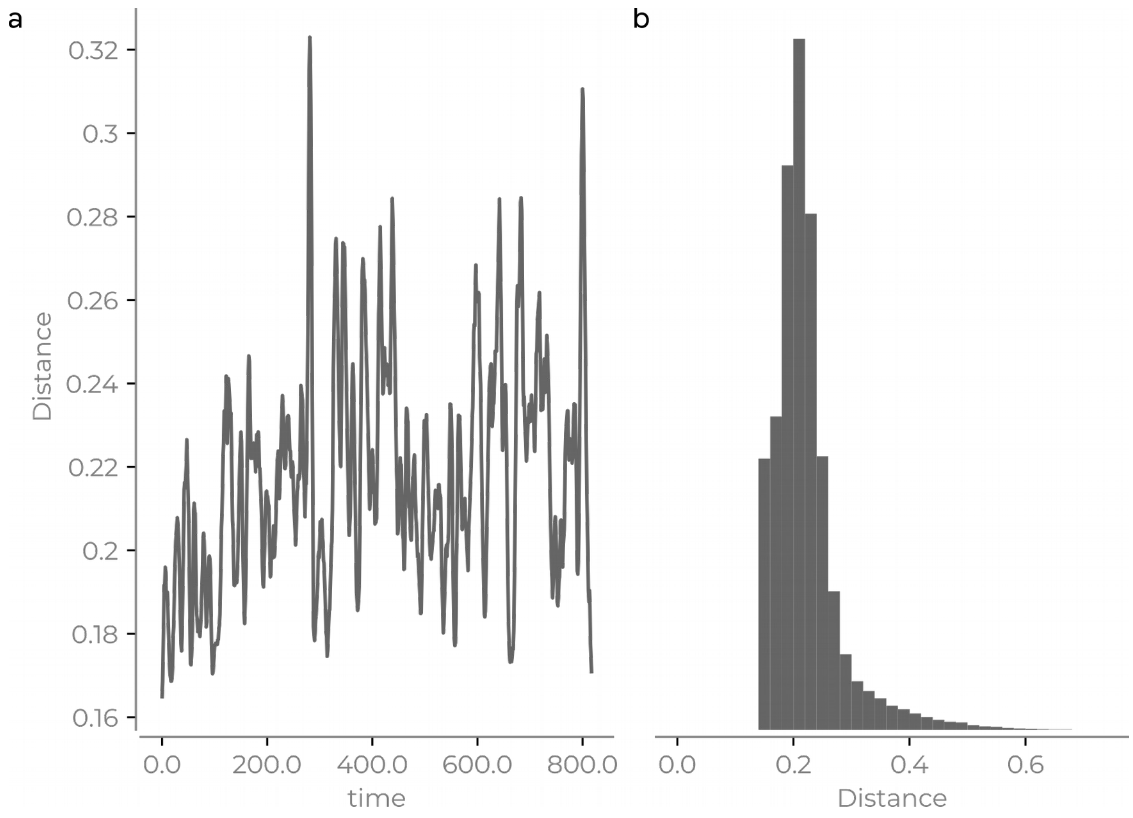


**Figure S4. RS temporal community fluctuations through time**. **a** Hamming distance for one example session/subject from the MSC dataset showing between the binary connectivity matrix based on the static template and the TCTC communities. **b** Distribution of hamming distance for all subjects and sessions illustrating a heavy tailed distribution.

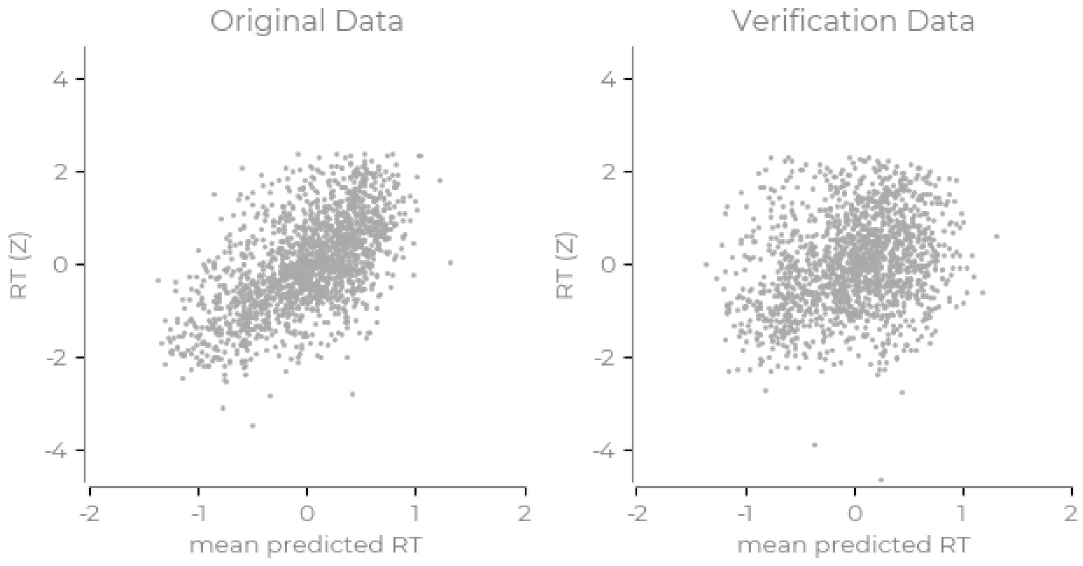


**Figure S5. Verification of RT model**. Left is the average predictive posterior samples vs the corresponding outcome variable on the original data. Right is the average predictive posterior samples vs the verification data.

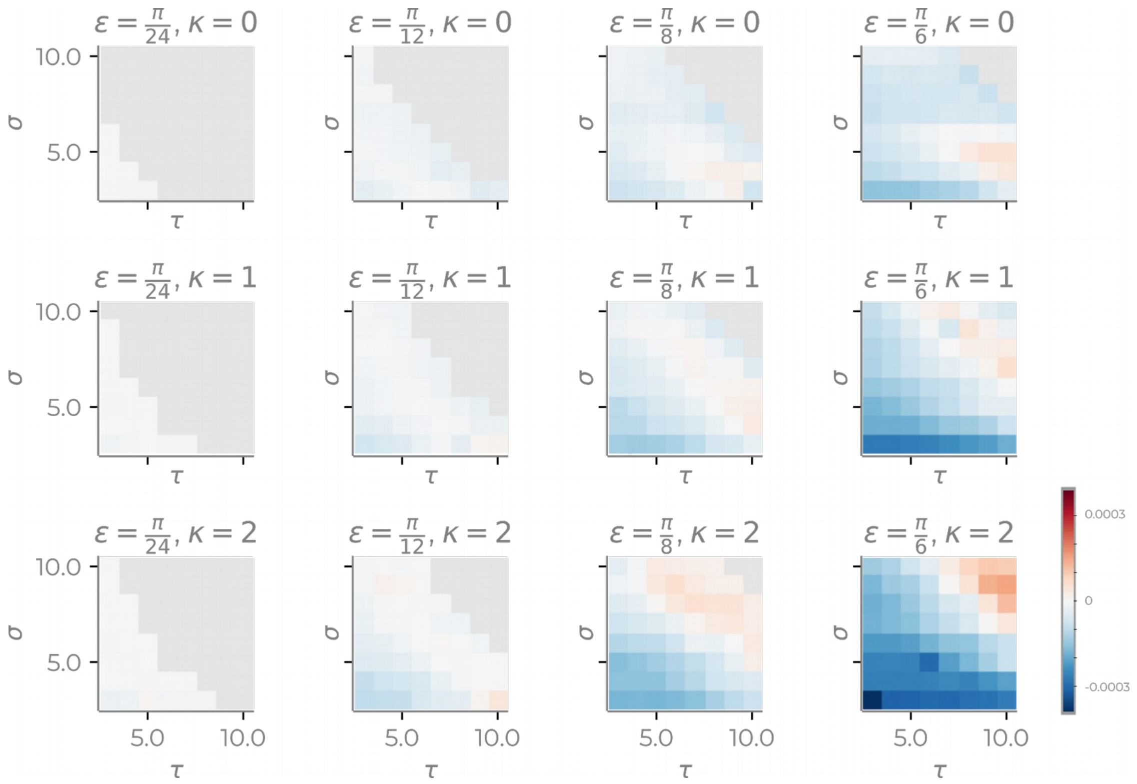


**Figure S6. Grid search optimisation when changing the 4 different hyper-parameters when contrasting 2-back vs 0-back**. Negative values indicate when the 2-back blocks were more dissimilar to 0-back blocks.

| | loo | p_loo | d_loo | weight | se | dse |
|---|---|---|---|---|---|---|
| **M-tcomb** | 4,641.79 | 155.34 | 0.00 | 0.61 | 63.04 | 0.00 |
| **M-t2** | 4,644.98 | 156.38 | 3.19 | 0.25 | 63.11 | 5.50 |
| **M-PC1-t2** | 4,646.93 | 152.98 | 5.14 | 0.01 | 62.86 | 5.77 |
| **M-t1** | 4,647.52 | 156.42 | 5.73 | 0.00 | 63.06 | 5.69 |
| **M-t3** | 4,648.77 | 156.33 | 6.98 | 0.00 | 63.13 | 6.02 |
| **M-t4** | 4,650.04 | 156.27 | 8.25 | 0.00 | 63.09 | 6.43 |
| **M-PC2-t6** | 4,650.29 | 153.70 | 8.50 | 0.13 | 62.83 | 7.18 |
| **M-PC4-t1** | 4,650.62 | 153.85 | 8.83 | 0.00 | 63.05 | 7.22 |
| **M-t0** | 4,650.86 | 156.30 | 9.07 | 0.00 | 62.96 | 6.54 |
| **M-t6** | 4,653.04 | 156.45 | 11.25 | 0.00 | 62.76 | 7.52 |
| **M-t5** | 4,653.41 | 156.36 | 11.62 | 0.00 | 62.92 | 7.14 |
| **M-t9** | 4,654.86 | 156.36 | 13.08 | 0.00 | 62.96 | 8.21 |
| **M-t10** | 4,655.03 | 156.84 | 13.24 | 0.00 | 63.10 | 8.65 |
| **M-t8** | 4,655.99 | 156.42 | 14.20 | 0.00 | 62.73 | 7.79 |
| **M-t7** | 4,657.17 | 156.69 | 15.38 | 0.00 | 62.72 | 7.20 |

**Table S1** LOO values for all models regarding reaction times. Explanation of model names: *M-tcomb* is all PC components from multiple time-points that are marked with a * in Figure 5. *M-PCX-tY* are models that are one PC component X at time-point Y. These models are the PC components marked with a * in Figure 5. *M-tX* are models that contain all 5 PC components at time-point X.

| | loo | p_loo | d_loo | weight | se | dse |
|---|---|---|---|---|---|---|
| **M-tcomb** | 1,017.48 | 67.57 | 0.00 | 0.54 | 55.91 | 0.00 |
| **M-PC2-t1** | 1,017.97 | 65.89 | 0.49 | 0.45 | 55.91 | 5.24 |
| **M-t0** | 1,024.46 | 70.16 | 6.98 | 0.00 | 56.43 | 4.82 |
| **M-t1** | 1,024.92 | 70.43 | 7.44 | 0.00 | 56.54 | 4.80 |
| **M-t2** | 1,026.34 | 68.72 | 8.86 | 0.00 | 56.46 | 4.70 |
| **M-PC4-t1** | 1,029.34 | 60.69 | 11.86 | 0.00 | 56.44 | 6.91 |
| **M-t3** | 1,029.71 | 68.19 | 12.23 | 0.00 | 56.59 | 5.40 |
| **M-PC4-t5** | 1,029.74 | 60.98 | 12.26 | 0.00 | 56.42 | 7.44 |
| **M-t4** | 1,032.82 | 67.45 | 15.34 | 0.00 | 56.65 | 5.86 |
| **M-t5** | 1,034.77 | 67.41 | 17.29 | 0.00 | 56.82 | 6.52 |
| **M-t6** | 1,041.31 | 68.21 | 23.83 | 0.00 | 57.24 | 8.13 |
| **M-t9** | 1,046.52 | 67.97 | 29.04 | 0.00 | 57.67 | 9.45 |
| **M-t8** | 1,046.97 | 68.09 | 29.49 | 0.00 | 57.68 | 9.55 |
| **M-t7** | 1,047.95 | 68.08 | 30.47 | 0.00 | 57.73 | 9.86 |
| **M-t10** | 1,051.26 | 70.09 | 33.78 | 0.00 | 58.03 | 10.38 |

**Table S2** LOO values for all models regarding accuracy. Explanation of model names: *M-tcomb* is all PC components from multiple time-points that are marked with a * in Figure 5. *M-PCX-tY* are models that are one PC component X at time-point Y. These models are the PC components marked with a * in Figure 5. *M-tX* are models that contain all 5 PC components at time-point X.

| | loo | p_loo | d_loo | weight | se | dse |
|---|---|---|---|---|---|---|
| M-tcomb | 523.04 | 29.93 | 0.00 | 0.96 | 49.95 | 0.00 |
| M-PC1-t2 | 525.41 | 28.80 | 2.37 | 0.00 | 50.12 | 2.69 |
| M-PC5-t10 | 528.18 | 28.59 | 5.14 | 0.04 | 50.19 | 4.85 |
| M-t0 | 530.99 | 33.97 | 7.95 | 0.00 | 50.51 | 4.79 |
| M-t1 | 531.86 | 34.45 | 8.82 | 0.00 | 50.65 | 5.10 |
| M-t2 | 534.31 | 33.36 | 11.27 | 0.00 | 50.96 | 5.64 |
| M-t3 | 536.73 | 32.27 | 13.69 | 0.00 | 51.30 | 5.94 |
| M-t4 | 537.85 | 30.48 | 14.82 | 0.00 | 51.35 | 6.06 |
| M-t5 | 539.28 | 30.44 | 16.25 | 0.00 | 51.58 | 6.29 |
| M-t6 | 542.19 | 30.24 | 19.16 | 0.00 | 51.75 | 6.50 |
| M-t7 | 543.42 | 27.95 | 20.38 | 0.00 | 51.74 | 6.64 |
| M-t8 | 543.71 | 32.18 | 20.68 | 0.00 | 51.75 | 6.74 |
| M-t9 | 547.70 | 27.66 | 24.67 | 0.00 | 52.20 | 7.62 |
| M-t10 | 548.55 | 33.55 | 25.51 | 0.00 | 52.23 | 7.84 |

**Table S3** LOO values for all models regarding response. Explanation of model names: M-*tcomb* is all PC components from multiple time-points that are marked with a * in Figure 5. *M-PCX-tY* are models that are one PC component X at time-point Y. These models are the PC components marked with a * in Figure 5. *M-tX* are models that contain all 5 PC components at time-point X.

| PC | Behaviour | Median of posterior | Time | 90% CI | % posterior above/below 0 |
|---|---|---|---|---|---|
| PC1 | Reaction time | 0.086 | 2 | [0.043, 0.13] | 99.93 |
| PC2 | Reaction time | 0.043 | 6 | [0.0047, 0.082] | 96.77 |
| PC4 | Reaction time | -0.046 | 1 | [-0.085, -0.0076] | 97.56 |
| PC2 | Accuracy | 0.30 | 1 | [0.14, 0.47] | 99.86 |
| PC4 | Accuracy | -0.20 | 1 | [-0.35, -0.045] | 98.47 |
| PC4 | Accuracy | -0.18 | 5 | [-0.33, -0.026] | 97.23 |
| PC1 | Response | 0.31 | 2 | [0.047, 0.61] | 97.50 |
| PC5 | Response | -0.30 | 10 | [-0.51, -0.093] | 99.05 |

**Table S4**: Description from creating new models combining credible features from __Figure 5cde__ where peaks of the median posterior where 90% of the credible interval were above/below zero. Results describe posterior distributions in Figure 6.
